# Development of visual cortex in human neonates are selectively modified by postnatal experience

**DOI:** 10.1101/2022.03.16.484671

**Authors:** Mingyang Li, Tingting Liu, Xinyi Xu, Qingqing Wen, Zhiyong Zhao, Xixi Dang, Yi Zhang, Dan Wu

**Affiliations:** Key Laboratory for Biomedical Engineering of Ministry of Education, Department of Biomedical Engineering, College of Biomedical Engineering & Instrument Science, Zhejiang University, Yuquan Campus, Hangzhou 310027, China; Department of Psychology, Zhejiang Sci-Tech University, Hangzhou 310018, China

**Keywords:** visual cortex, development, neonates, MRI, experience-dependent plasticity

## Abstract

Experience-dependent cortical plasticity is a pivotal process of human brain development and essential for the formation of most cognitive functions. Although studies found that early visual experience could influence the endogenous development of visual cortex in animals, little is known about such impact on human infants. Using the multi-modal MRI data from developing human connectome project, we revealed the early structural and functional maps in the ventral visual cortex and their development across the first month of age. Particularly, we found the postnatal experience could modulate both the cortical morphology in ventral visual cortex and the functional circuit between bilateral primary visual cortices. But the cortical myelination and overall functional circuits of ventral cortex, particularly that of the high-order visual cortex, developed without significant influence of postnatal experience in such early period. These experience-dependent cortical properties were further validated in the preterm-born infants who have longer postnatal time but immature cortical development at birth. The results confirmed that the development of cortical thickness was dominated by the postnatal experience but the functional circuit might be determined by the overall maturity of visual cortex. These data suggest in human newborns that early postnatal experience shapes the structural and functional development of the visual cortex in selective and organized pattern.

## Introduction

A fundamental question in neuroscience is about the role of experience in neurodevelopment (Arcaro and Livingstone, 2021; Barlow, 1975; Holtmaat and Svoboda, 2009; Nithianantharajah and Hannan, 2006). Vision, given its ecological universality and importance across species, has long been taken as a representative modality to investigate such question (Barlow, 1975; Crair et al., 1998; Gödecke and Bonhoeffer, 1996; Li et al., 2008, 2006; Roy et al., 2020), and a framework consisting of two distinct phases was proposed to describe the development of visual cortex (Barlow, 1975; Li et al., 2006; White and Fitzpatrick, 2007). This framework includes an early, experience-independent phase in which the basic layout of neural map is established, and a subsequently experience-dependent phase in which the visual experience refines and shapes the initial neural map. Recent studies further revealed the importance of visual experience during the early period of cortical development, suggesting the intricate interaction between early sensory experience and the endogenous mechanisms in neurodevelopment (Li et al., 2008, 2006; Roy et al., 2020). However, these studies were carried out on the model animals (e.g. cat and ferret) using electrophysiological techniques. Due to the lack of non-invasive methods to probe the normal human infant brain, whether early postnatal experience have influence on the development of visual cortex in human infants remains unclear.

Neuroimaging techniques, especially magnetic resonance imaging (MRI), provided an ideal tool to noninvasively measure both the brain structure and function, which however, remains challenging for the infant brain due to subject motion, demand for resolution, and difficulty in patient recruitment (Cordero-Grande et al., 2018). Owing to the recent technical advances in both images acquisition and processing methods (Hughes et al., 2017), the developing human connectome project (dHCP; http://www.developingconnectome.org/) was initiated to investigate the early structural and functional development of human cerebral cortex. Herein we collected a dataset from the dHCP that included multi-modal MRI images of 783 neonatal subjects to address above question. The different time interval between gestational age (GA) at birth and the postmenstrual age (PMA) at scan indicated the difference in the postnatal experience across neonates, which may lead to the individual variation of the visual experience. Thus this time window provides us the opportunity to investigate the contribution of early visual experience versus endogenous neurodevelopment on the development of visual cortex in human newborns.

The main purpose of present study is to describe the early structural and functional development of the ventral visual cortex in human newborns and estimate the contribution of postnatal experience on this process. Particularly, the cortical thickness (CT, Lyall et al., 2015; Sowell et al., 2004) and T1w/T2w-based cortical myelination (CM, Glasser and van Essen, 2011; Soun et al., 2017) data from dHCP was used to investigate the development of cortical morphology and microstructure, respectively. The resting-state functional MRI (r-fMRI, Fitzgibbon et al., 2020) data was used to evaluate the development of cortical circuit based on function connectivity between cortical areas. We focused on the primary visual cortex (V1) and higher-level visual cortex, namely, the ventral occipital temporal cortex (VOTC, Bi et al., 2016; Grill-Spector and Weiner, 2014). Area V1 was selected because this region is the most basic region for visual processing and probably is the most experience-dependent area during early development. The VOTC contains function-specific regions for biologically important categories such as faces (Kanwisher et al., 1998), bodies (Downing et al., 2001) and scenes (Epstein Russell and Kanwisher, 1998), making this region a critical component in the ventral pathway of visual processing (Bi et al., 2016; Grill-Spector and Malach, 2004; Grill-Spector and Weiner, 2014; Kanwisher, 2010), and whether early visual experience has influence on this region is an interesting question.

Previous studies have described the developmental trajectory of CT and myelination in human infants, and found a general increasing tread for both two measurements with PMA in whole brain (Bozek et al., 2018; Fenchel et al., 2020). But the contribution of postnatal experience to the development were not clarified because PMA reflects both prenatal endogenous effect and postnatal experience. Using r-fMRI, studies found that the primate newborns with a few days of age already had a proto-organization in visual system (Arcaro and Livingstone, 2017) and human infants with a few weeks of age showed category-specific network (Kamps et al., 2020). However, the small sample size of those studies might undermine reliability of the results given the well-known instability of r-fMRI signals (Poldrack et al., 2017). Moreover, those studies did not fully control the visual experience in the first few weeks of the subjects, thus cannot give a clear conclusion whether the innate functional connectivity is unrelated to postnatal visual experience.

In the present study, we first decipher endogenous and external effects on early structural development in human visual cortex by examining the correlation between morphology (CT) and microstructure (CM) with respect to PMA and postnatal time of infants. We found that both two structural measurements significantly increased from 37 to 45 weeks of PMA in ventral visual cortex, but they were differently influenced by postnatal experience. Using r-fMRI, we characterized the innate organization of the ventral cortex in the infants as early as one day of life and investigated the effect of postnatal time on functional networks of V1 and VOTC areas. Finally, we evaluated the influence of postnatal experience and innate maturity on visual cortex development by comparing the cortical measurements in the term- and preterm-born babies, as the latter group had immature cortical development at birth but longer postnatal time compared to former group with equivalent PMA.

## Results

### General development of cortical thickness and cortical myelination in human infants

The present study included 407 neonates (187 females, PMA = 41.11 ± 1.73 weeks at scan) from a total dataset of 783 subjects after excluding data that did not pass quality control or did not satisfy inclusion criteria (STAR Methods), and 355 of them were term-born (PMA = 41.14 ± 1.70, range from 37.43 to 44.71 weeks at scan; Figure 1a). The general trend and spatial variation of visual cortex development was described within an anatomical mask of ventral cortex, which was then segmented into 34 regions-of-interest (ROIs) per hemisphere according to the HCP-MMP atlas (Glasser et al., 2016), including the early visual cortex (e.g. V1 and V2), higher-level visual cortex (e.g. VOTC) and anterior part of temporal cortex (Figure S1 and Table S1). In general, the CT and CM of the ventral cortex significantly increased between 37 to 45 weeks of PMA (*rs* = 0.40 to 0.49, *ps* < 10^−9^; Figure 1b-d). We found distinct spatial variation in the developmental pattern of the 34 ROIs in ventral cortex along different direction (e.g. posterior-anterior and medial-lateral directions; Figure S2a). Both CT and CM show higher correlation with PMA in the posterior than anterior region, and higher correlation in the medial than lateral part within the anatomical mask (Figure 2a and Figure S2b-c).

**Fig 1.**
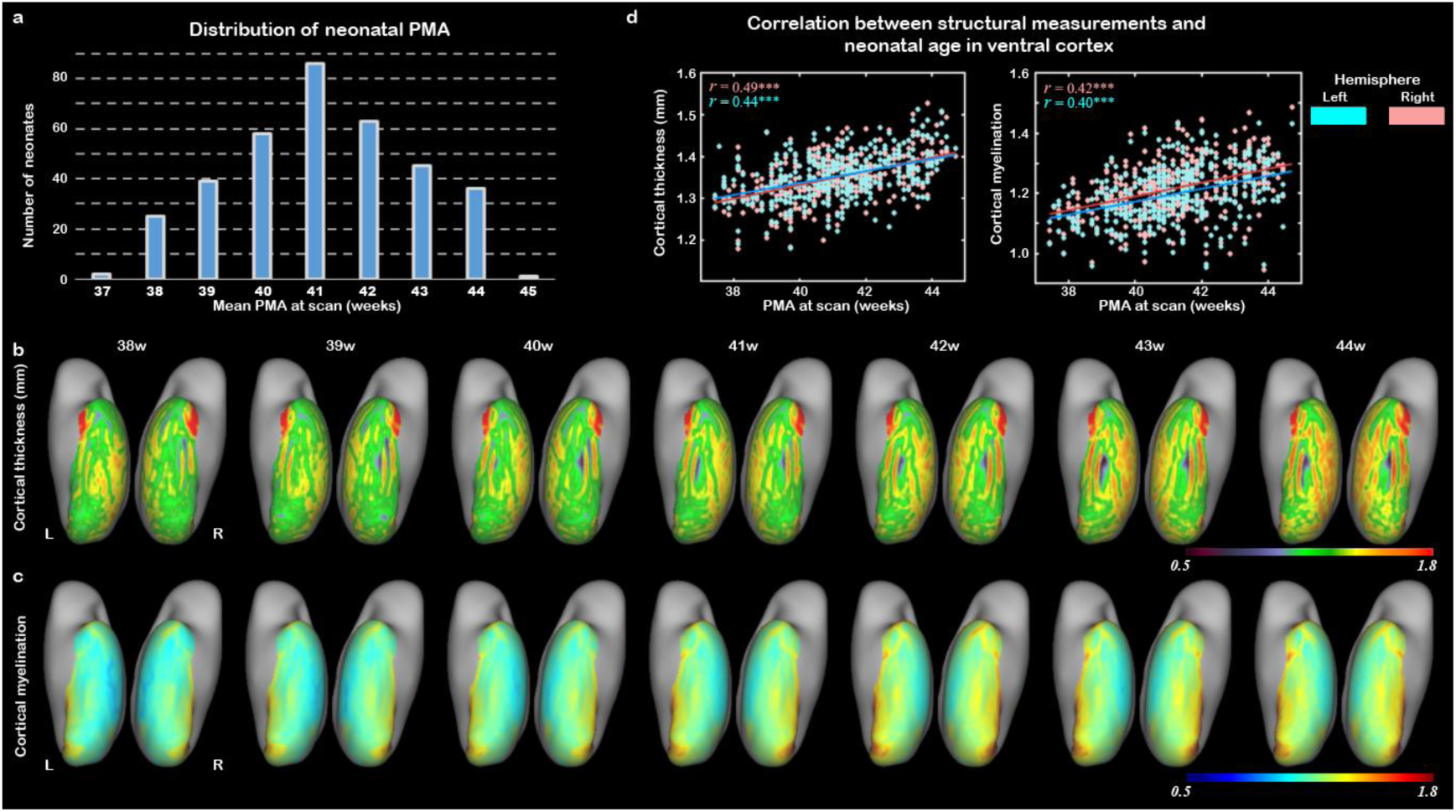
The development of cortical structural properties in human newborns. (a) The distribution of neonatal postmenstrual age (PMA) at scan in the study population; (b) The averaged cortical thickness (CT) and (c) the averaged cortical myelination (CM) from 38 to 44 weeks of PMA in the ventral cortex; (d) The correlation between CT/CM and PMA in right (peach) and left (blue) ventral cortex (*** *p* < 0.001).

**Fig 2.**
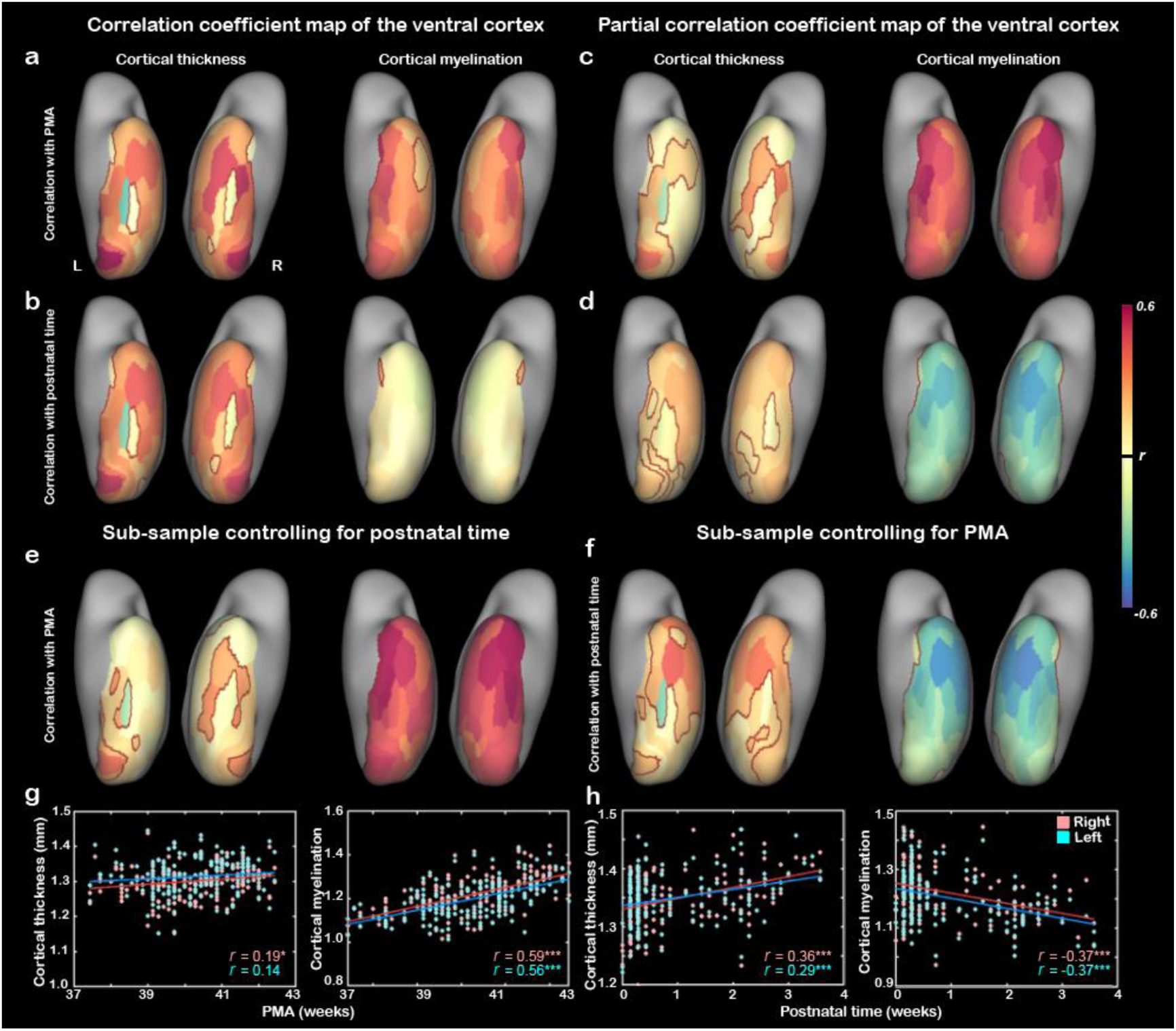
The correlation maps between cortical thickness/cortical myelination and PMA (a) or postnatal age (b) in ventral cortex. Partial correlation maps between cortical structural indices and PMA controlling for the postnatal age (c) or between the cortical measurement and postnatal time controlling for PMA (d). (e-h) Validation analysis using two sub-samples: the first sub-sample included infants who underwent the scans within 3 days after birth to control for the postnatal time (e, g) and the second sub-sample included infants whose PMA were ranged from 40 to 42 weeks to control for the chronological age (f, h). The scatter plot illustrates the relationship between CT/CM in ventral cortex and PMA in the first sub-sample (g) or between these measurements and postnatal time in the second sub-sample (h). Note: correlation coefficients in the areas with the maroon contours were significant after FDR correlation (FDR *q* < 0.05); PMA = postmenstrual age at scan; * *p* < 0.05 and *** *p* < 0.001 by Pearson correlation.

### Contribution of postnatal time on the development of cortical thickness and cortical myelination

Although both two structural measurements in ventral cortex increased with PMA, the influence of postnatal time on them were different. The postnatal time, defined as PMA-GA, was significantly correlated with CT (*rs* = 0.46 and 0.51 for right and left hemisphere, *ps* < 10^−9^; 61/68 ROIs showed significantly positive correlation; *rs* > 0.11, FDR corrected *q* < 0.05) but not CM (*rs* = 0.02 and 0.01, *ps* > 0.6; 2/68 ROIs showed significant effect) of ventral cortex (Fig 2b and Supplementary Fig 4a-b). Given the high correlation between postnatal time and PMA (*r* = 0.68, *p* < 10^−9^), partial correlation analysis was applied to regress out one effect from the other. The CT in part of ventral cortex (28/68 regions) were still significantly correlated with the PMA, but the explained variation (*R*^*2*^) were significantly decreased in all 68 ROIs (*t* = -8.83, *p* < 10^− 9^) after regressing out the effect of postnatal time (Fig 2c). The partial correlation between CT and postnatal time remain significant in most ROIs (51/68 regions) in the ventral cortex but became weaker after regressing out the effect of PMA (*t* = -5.58, *p* < 10^−5^; Fig 2d). Those results suggested that both postnatal experience and endogenous growth contributed the development of CT in the ventral cortex. In contrast, the correlation between CM and PMA in all ROIs even became stronger after controlling for the postnatal time (*t* = 11.34, *p* < 10^−9^; Fig 2c), and the correlation between CM and postnatal time altered to negative in all ROIs after controlling for PMA (Fig 2d). This seemly counterintuitive results might reflect the impact of prenatal time on the myelination because subjects who had longer postnatal time would have lower GA at birth at a given PMA. To test this assumption, we first applied correlation analysis between GA and cortical measurements, and found that both CT (*rs* = 0.16 and 0.14; *ps* < 0.05) and CM (*rs* = 0.53 and 0.51; *ps* < 10^−9^) showed significant correlation with the GA of the infants (Supplementary Fig 4d-e). To test whether GA at birth had stronger influence on CM than CT, we separated the infants into two groups, the first group contained the subjects born at relatively high GA (top 1/4) and the second group born at relatively low GA (low 1/4). We found significant interaction between prenatal time (high and low GA at birth) and the two structural measurements (See supplementary methods, the interactive effect *F*_inter_ = 81.2, *p* < 10^−9^; Supplementary Fig 4f), and the interactoin was more prominent in CM (simple effect: *t* = 10.98, *p* < 10^−9^) that in than CT (*t* = 2.07, *p* < 0.05). In contrast, the postnatal time showed inverse interaction in the two measurements (*F*_inter_ = 6.82, *p* < 0.01), in which the postnatal time have stronger influence on the CT than CM (Supplementary Fig 4c).

Furthermore, two sub-samples of the dataset were selected to validate above results. The first sub-sample included infants who underwent the scans within 3 days after birth (*n* = 173) and the second sub-sample included infants whose PMA were within a short range from 40 to 42 weeks (*n* = 163). If the postnatal experience does increase CT independent of PMA, a weak or insignificant correlation between CT and PMA would be expected in the first sub-sample who had very limited postnatal experience, but significant postnatal correlation would be found in the second sub-sample even though we controlled the PMA of infants. In the first sub-sample, CM showed strong correlation with PMA (*rs* = 0.59 and 0.56 for right and left hemisphere, *ps* < 10^−9^) but not the CT (*r* = 0.19 and 0.14, *p* < 0.02 and > 0.07; Fig 2e, g). In the second sub-sample, significantly positive correlation remained between CT and postnatal time in the ventral cortex (*r* = 0.36 and 0.29, *ps* < 0.001), but significantly negative correlation was found between CM and postnatal time (*r* = -0.37 and -0.37, *ps* < 10^−6^; Fig 2f,h). Those results supported above findings that the development of CM heavily depend on the endogenous mechanisms and longer prenatal time would be beneficial for the development of cortical myelination. In contrast, the development of CT in neonates was driven by both postnatal time and endogenous growth.

Next we focused on two typical visual including the V1 (V1 patch) and VOTC (15 patches) based on HCP-MMP atlas (Fig 3a and Supplementary Figure 1 and Table 1). Both regions showed significant increase of CT and CM with PMA (*rs* = 0.39 – 0.61; *ps* < 10^−9^; Fig 3b-c and Supplementary Fig 5). Similarly, only the development of CT was modulated by the postnatal time (*rs* = 0.26 – 0.52, *ps* < 10^−6^ for the correlation analysis; *rs* = 0.17 – 0.21, *ps* < 0.01 for the partial correlation analysis controlling for the PMA; Fig 3b-c). Moreover, CT in V1 showed faster thickening compared to VOTC with both PMA and postnatal age (*F* = 35.98 and 12.15, *p* < 0.001; Fig 3d-e). To validate the postnatal effect on the two visual cortical areas, we applied similar analysis in the two sub-samples defined above. We found that the increase of CT remained significant in V1 (*r* = 0.18 -0.40; *ps* < 0.05; Fig 3f-g) in above two sub-samples controlling for either postnatal time or PMA. But only the increase of CT in the right VOTC (*r* = 0.26 and 0.04; *p* < 0.01) kept significant in the two sub-samples (*r* = 0.02 and 0.04; *p* > 0.5 in the left VOTC; Fig 3f-g), indicating an asymmetrical effect of the development of the higher-level visual cortex between two hemispheres.

**Fig 3.**
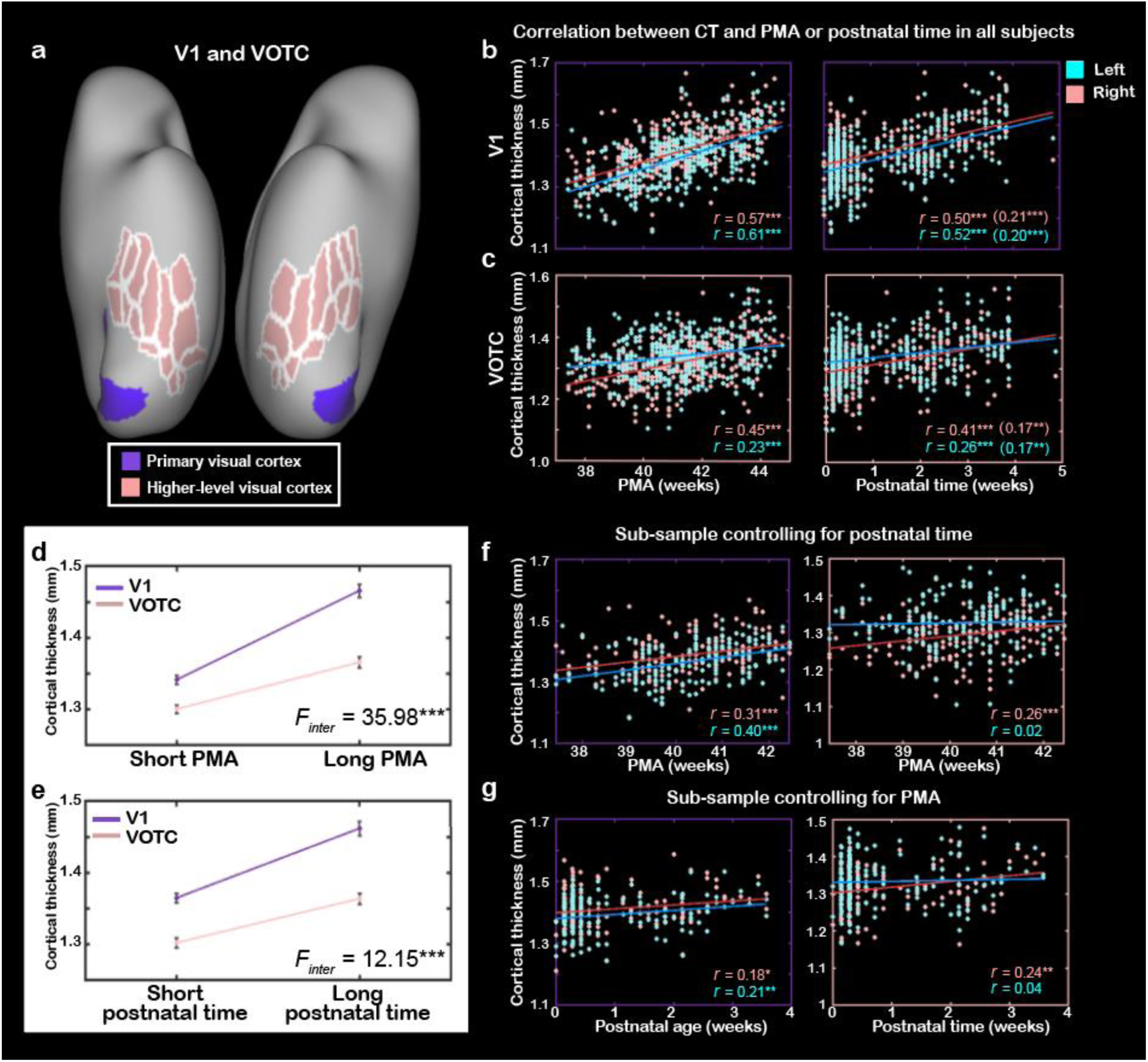
The development of cortical thickness (CT) in primary visual cortex and higher-level visual cortex in human newborns. (a) Definitions of primary visual cortex (V1) and higher-level visual cortex (VOTC). Correlation between cortical thickness and PMA or postnatal age in V1 (b) and VOTC (c), and the values in the bracket indicate the partial correlation between CT and postnatal time when controlling for PMA. The averaged CT in the short (e.g. top 25%) and long (e.g. last 25%) PMA (d) or postnatal age (e) groups. Validation analysis in the two sub-samples controlling for the postnatal time for subsample 1 (f) and the PMA for subsample 2 (g), respectively. Note: V1 = primary visual cortex; VOTC = ventral occipital temporal cortex; * *p* < 0.05; ** *p* < 0.01 and *** *p* < 0.001.

**Fig 4.**
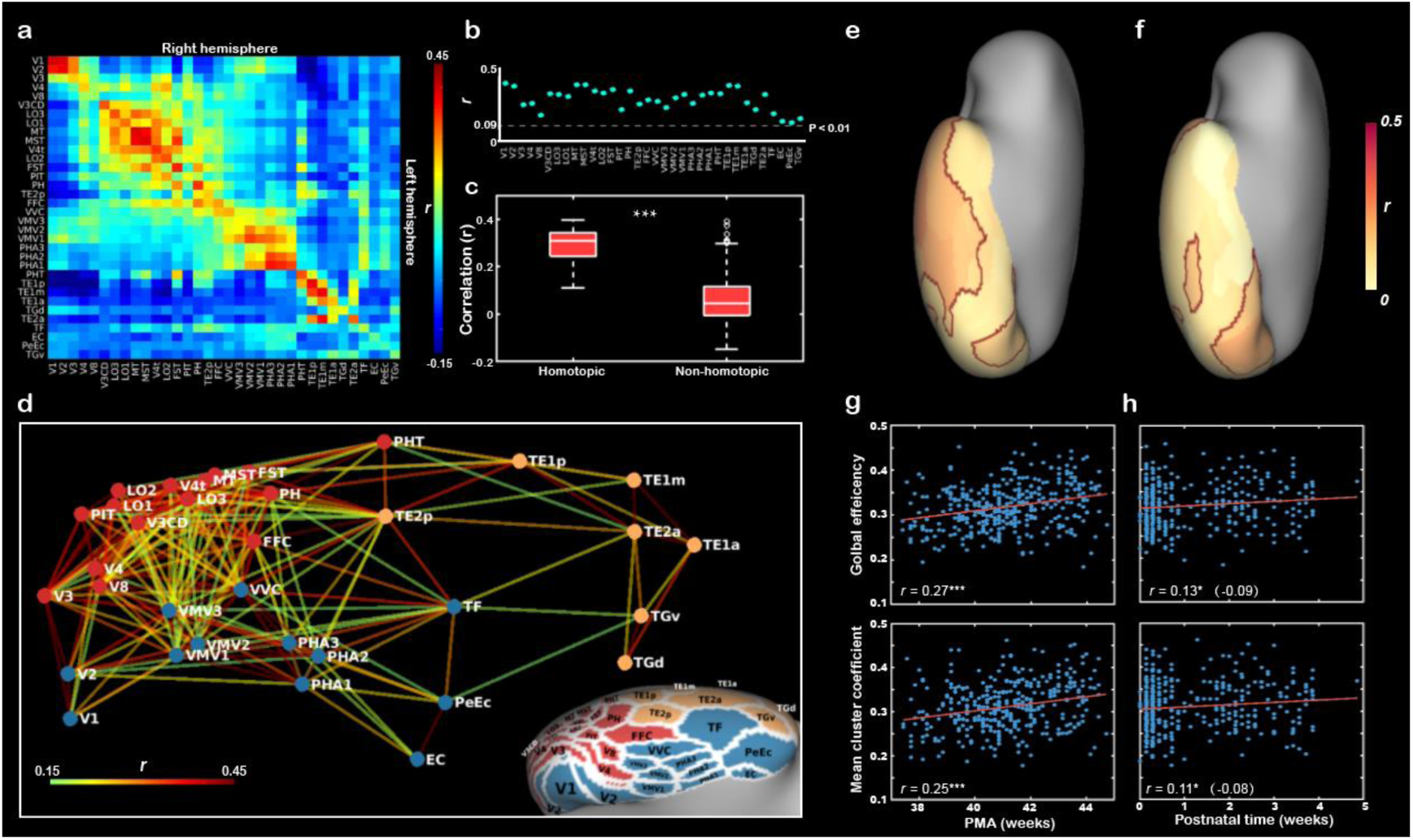
The innate functional organization of ventral cortex within 1 day after birth and its development in the first month of age in human newborns. (a) The pairwise correlation matrix describes the functional correlations among 34 ROIs across hemispheres in ventral cortex. (b) The scatter graph illustrates the correlation coefficients (*r*) between 34 pairs of bilateral homotopic areas, and the dash line indicates the corrected threshold (Bonferroni corrected *p* < 0.01). (c) The box graph depicts the comparison between homotopic and non-homotopic connections at birth. (d) Multidimensional scaling and community groups obtained from the pairwise ipsilateral connections between 34 ROIs in right ventral cortex and the corresponding projection on the cortical surface. The color of the nodes in (d) indicate the cluster identity in the community structure analysis and color of the lines connecting the nodes indicate the functional correlation (*r* value) between two nodes within the right hemisphere. The correlation coefficient maps between the homotopic correlation and PMA (e) or postnatal time (f) in 34 ROIs, the maroon contours indicated the significant regions after FDR correlation (FDR *q* < 0.05). The correlation between network measurements of the right hemisphere and PMA (g) or postnatal time (h). The values in the bracket indicate the partial correlation coefficients in (h) between the network properties and postnatal time when controlling for PMA. * *p* < 0.05, *** *p* < 0.001.

**Fig 5.**
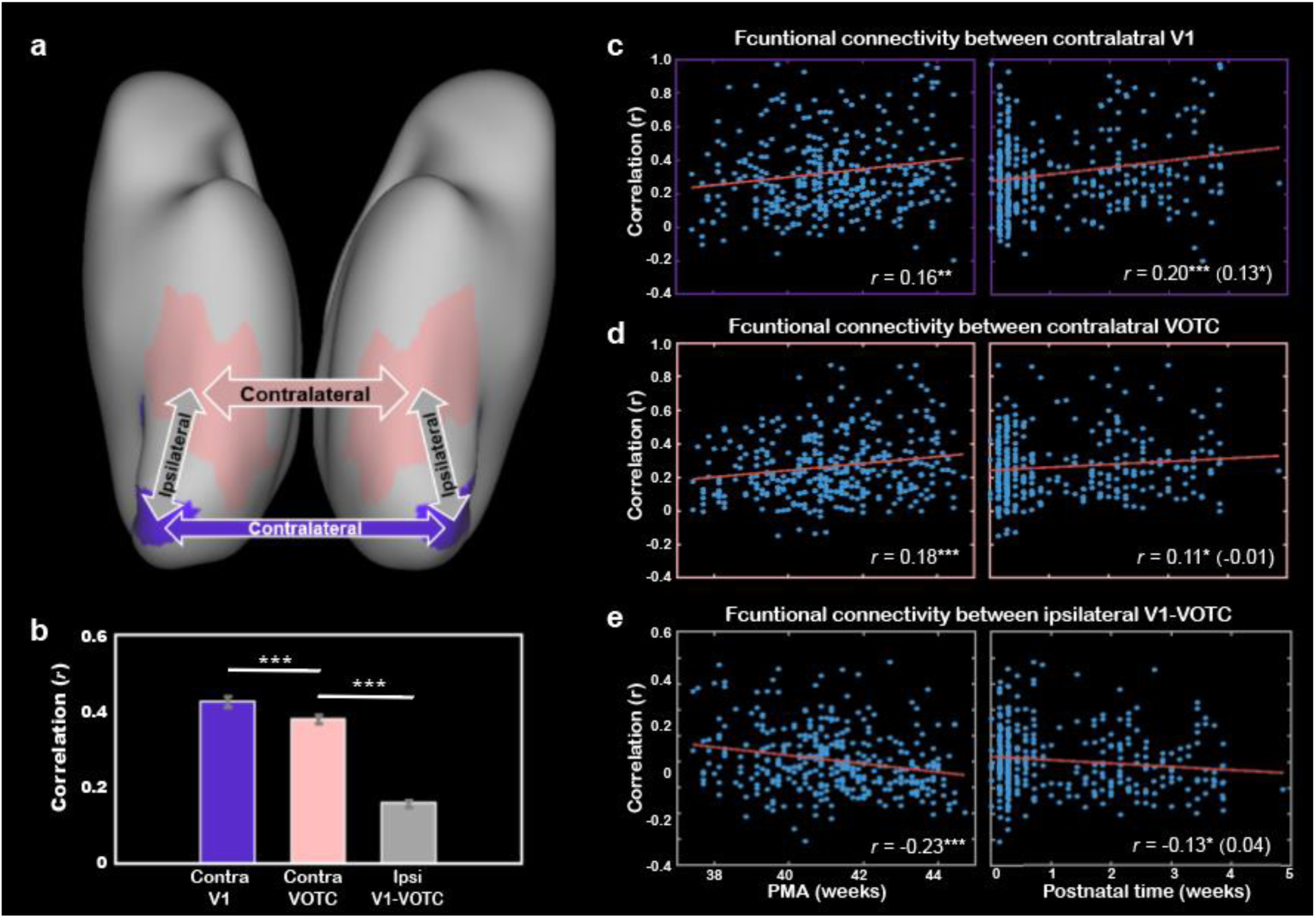
The development of the functional connections between bilateral primary visual cortex and higher-level visual cortex and their cross-correlations. (a) The connections of interest include the homotopic connection between bilateral V1 (purple), bilateral VOTC (peach) and the averaged ipsilateral connections between V1 and VOTC in each of the hemispheres (gray). (b) Comparison of the three types of connections in the infants within 1 day after birth (no postnatal experience). The correlation between PMA or postnatal age and bilateral V1 connection (c), bilateral VOTC connection (d) and ipsilateral connection between V1 and VOTC (e). The values in the bracket indicate the partial correlation coefficient controlling for PMA of infants. Note: PMA = postmenstrual age; Contra = contralateral; Ipsi = ipsilateral; * *p* < 0.05; ** *p* < 0.01 and *** *p* < 0.001.

### Innate functional organization of ventral cortex in human infants

Beyond the relation between cortical structural development and postnatal and prenatal influences, we further asked how the connectivity between the different visual sub-regions change during early development. To do so, we first estimate the initial functional connectivity pattern in ventral cortex without the influence of postnatal visual experience in a sub-sample of subjects who underwent the scans within the first day after birth (*n* = 73). Homotopic correlation was calculated to reflect the functional distinction in neonatal ventral cortex (Arcaro and Livingstone, 2017; Vincent et al., 2007). The homotopic connections in all ROIs of ventral cortex were significant (mean *rs* = 0.11– 0.40; Bonferroni corrected *ps* < 0.01; Fig 4a-b), and were significantly higher than non-homotopic connections (*t* = 13.68, *p* < 10^−9^; Fig 4c), suggesting that arealization of ventral cortex already existed at birth. In addition, multidimensional scaling (MDS) analysis of the correlations between 34 ROIs within the right hemisphere revealed the functional relationship among those areas in ventral cortex (Fig 4d). Using community structure analysis on the network matrix (threshold *r* = 0.15, *p* < 10^−8^;Arcaro and Livingstone, 2017), we further partitioned these areas into 3 groups including a lateral cluster, a medial cluster, and an anterior-lateral temporal cluster (Fig 4d). Similar results were found with different threshold (e.g. *r* = 0.10, 0.15 and 0.20) and in the left hemisphere (Supplementary Fig 6). Taken together, those results suggested that the proto-organization of ventral visual cortex was formed before acquiring any higher-level visual experience.

**Fig 6.**
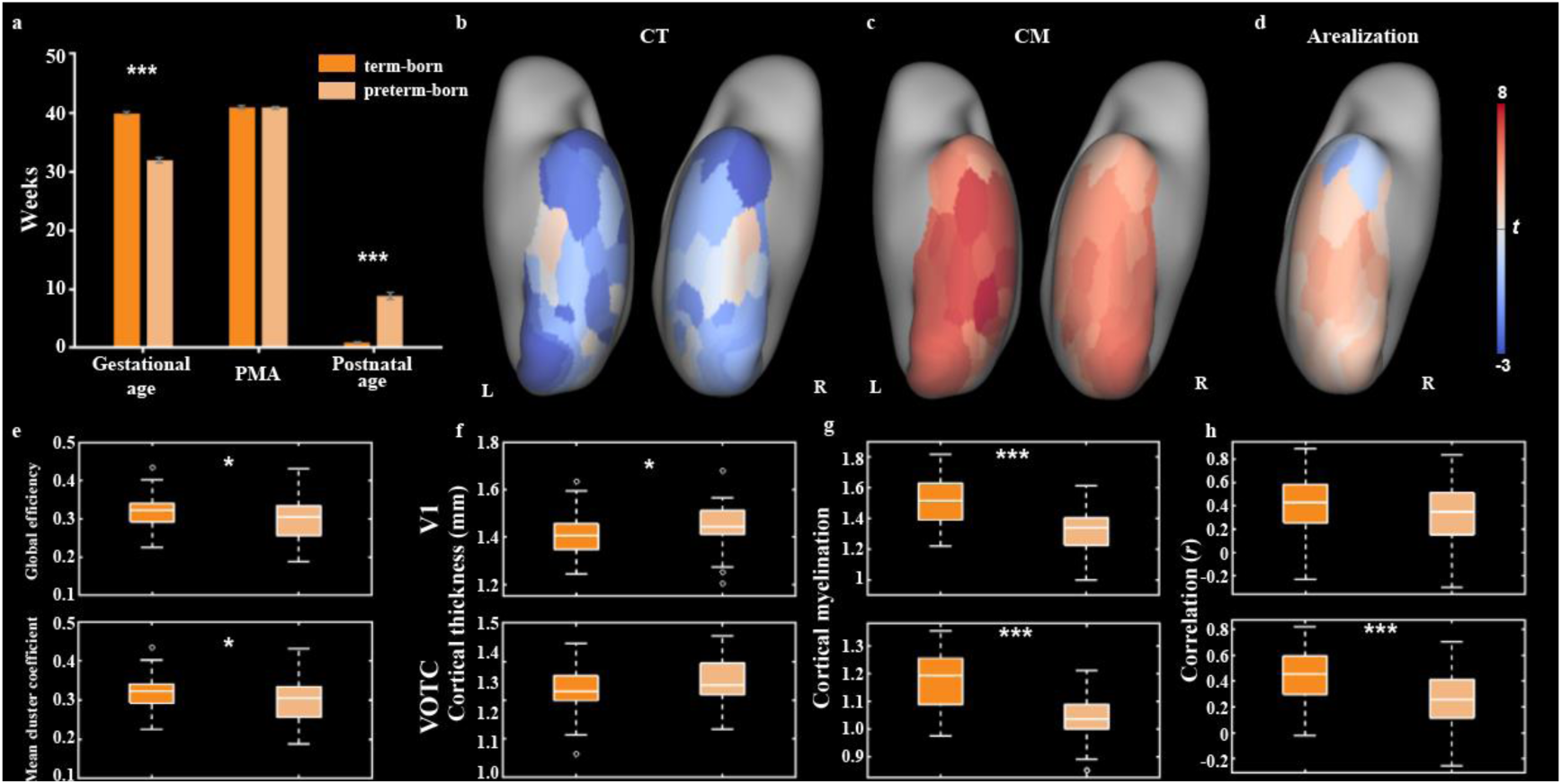
(a) The comparison of gestational age, postmenstrual age and postnatal age between term-born and preterm-born infants. (b)-(d) The t maps of cortical thickness (CT), cortical myelination (CM) and arealization between two groups (term -preterm) in ventral visual cortex. (e)-(h) The box plots of structural and functional measurements between two groups, including the global coefficient and mean cluster coefficient of the ventral network (e), cortical thickness in V1 and VOTC (f), cortical myelination in V1 and VOTC (g) and functional connectivity between contralateral V1 and VOTC (h). * *p* < 0.05; ** *p* < 0.01 and *** *p* < 0.001 by two sample *t* test.

### Development of functional connections in the ventral visual cortex

Whether those functional properties in ventral visual cortex develop in such early stage of human infants? If so, whether postnatal experience could modulate this course? For the arealizaiton in the 34 ROIs, primary visual cortex and lateral-anterior regions showed significant increase with PMA (16/34 regions, *rs* = 0.12 – 0.27; FDR *q* < 0.05; Fig 4e), and the arealization in primary visual cortex, occipital-temporal and anterior temporal patches were significantly related to postnatal time (6/34 patches, *rs* = 0.14 – 0.22; FDR *q* < 0.05; Fig 4f). Among the six ROIs, only the arealization in early visual cortexes (e.g. V1 and V2) showed significant correlation with postnatal time in the partial correlation analysis controlling for the PMA (*rs* = 0.17 and 0.14, *ps* < 0.01; Supplementary Fig 8). For the entire ventral cortex, we applied MDS and community structure analysis in each PMA week from 38–44 weeks across all term-born infants (n = 355), and found similar three-cluster network structure in the infants with different PMA (except for 42 weeks; Supplementary Fig 7). Then we used two typical network measurements including global efficiency and mean cluster coefficient to quantify the development of the functional network in ventral cortex. Both two measurements showed significantly increase with PMA between 37 and 44 weeks (*rs* = 0.27 and 0.25, *ps* < 10^−5^) and with postnatal time between 0 and 5 weeks (*rs* = 0.11 and 0.13, *ps* < 0.05), but the postnatal effects were not significant after we controlling for the PMA (*rs* = -0.09 and -0.08, *ps* > 0.1; Fig 4g-h).

In terms of the functional connections within the V1 and VOTC (Fig 5a), the homotopic correlation between contralateral V1’s (*r* = 0.43 ± 0.25) were significantly higher than the correlation between VOTCs (*r* = 0.38 ± 0.22; *t* = 4.29, *p* < 10^−4^), which were both higher than the connections between V1-VOTC within the same hemisphere (*r* = 0.16 ± 0.19; *ts* = 15.74 and 13.49, *ps* < 10^−9^; Fig 5b) across all the term-born infants. Furthermore, the homotopic correlations of both V1 and VOTC showed significant increase with PMA (*rs* = 0.16 and 0.18, *ps* < 0.01) but the connection between V1 and VOTC showed significant decrease with PMA (*r* = -0.23, *p* < 10^−4^). Moreover, the homotopic correlation of V1 was significantly modulated by the postnatal time with or without controlling for PMA (*rs* = 0.20 and 0.13, *ps* < 0.05; Fig 5c). Other connections were not significantly changed with postnatal time when controlling the effect of PMA (*ps* > 0.5; Fig 5d-e). To validate the postnatal effect in V1, we applied similar analysis in above two sub-sample of infants as described above. The homotopic connection of V1 showed no significant change with PMA (*r* = -0.04, *p* > 0.5) in the first sub-sample who had similar postnatal time, but showed significant increase with the postnatal time (*r* = 0.25, *p* < 0.002) in the second sub-sample who had similar PMA (Supplementary Fig 9).

### Comparison between structural and functional properties in ventral visual cortex of term- and preterm-born infants

Relative to the term-born infants, preterm-born infants have longer postnatal experience with relatively immature development at birth, and thus the comparison between them may help us to understand the influences of early experience and innate maturity on cortical development. Considering the unbalanced sample size between two groups (*n* = 355 vs. 52), we selected a sub-group in the term-born infants with equal sample size and similar PMA to the preterm-born neonates (Fig 6a). For the structural measurements, 56 of 68 ROIs showed significant lower CT in the term-born than preterm-born infants (*ts* = -4.26 to -2.71; FDR *q* < 0.05; Fig 6b). Particularly, the term-born infants showed significantly higher CT in the area V1 (*t* = -2.55, *p* < 0.02) but not VOTC (*t* = -1.24, *p* > 0.2; Fig 6f). In addition, all 68 patches showed significantly higher CM in the term-born than preterm-born infants (*ts* = 2.02 to 7.91, FDR *q* < 0.05; Fig 6c), and such trend were consistent in both area V1 and VOTC (*ts* = 6.65 and 7.54, *ps* < 10^−8^; Fig 6g). Those results supported the previous findings that early thickening of CT in neonates could be modulated by postnatal time but CM was heavily depending on the GA at birth.

For the functional organization of ventral cortex, two group showed similar three-community structure except that some areas (e.g. V1 and V2; Supplementary Fig 10) were included in other adjacent community in the preterm-born group. Significantly different network measures were found between two groups, e.g., the term-born infants had higher global efficiency and mean cluster coefficient than preterm-born infants (*t* = 2.22 and 2.43; *ps* < 0.05; Fig 6e), suggesting higher efficiency of information transmission and higher differentiation of cortical function in the ventral cortex in term-born infants. Interestingly, compared to preterm-born infants, the arealization in the term-born infants was higher in posterior and middle part of ventral cortex but lower in the anterior temporal areas (Fig 6d). Particularly, the preterm-born infants showed lower arealization in the VOTC (*t* = 3.75, *p* < 0.001) but not V1 (*t* = 1.60, *p* > 0.1) than term-born neonates (Fig 6h), suggesting that the connections between bilateral visual cortex was also influenced by the developmental maturity.

## Discussion

Experience-dependent plasticity is one of the most striking features of human brain and also the foundation of acquired cognitive ability. Utilizing the large datasets of neonatal multi-modal MRI images from dHCP, we reported the early structural and functional maps in the ventral visual cortex and their development across the first month of age, focusing on the contribution of postnatal experience to the cortical development in this early period. One the one hand, we found that the cortical myelination and overall functional connectivity of ventral cortex developed significantly with the PMA but was not directly influenced by postnatal time, suggesting the endogenous stability in the development of ventral visual cortex in human newborns. On the other hand, we found that postnatal experience could accelerate the cortical thickening and modify specific functional circuits (e.g. homotopic functional connections between the bilateral visual cortex) in the visual cortex, suggesting that the postnatal experience could selectively modify the structure and function of visual cortex as early as the first few weeks after birth. Furthermore, the influence of postnatal experience on the cortical development was also supported by the comparison between the preterm- and term-born infants; the former experienced longer postnatal stimuli at equivalent PMA. Taken together, those results suggested that the development of human visual cortex does not strictly follow the two-stage hypothesis (Barlow, 1975; Li et al., 2006; White and Fitzpatrick, 2007), as the cortical properties are modulated by postnatal experience even within the first month of life in both term- and preterm-born infants.

Similar to the previous findings in human infants (Bozek et al., 2018; Fenchel et al., 2020), both the CT and CM in ventral cortex increased between 37 and 44 weeks of PMA in our study. We went one step to investigate the different mechanism underlying these trends, and found the development of CT was considerably modulated by the postnatal experience while the CM was heavily influenced by prenatal duration. CT is related to the synaptogenesis and synaptic pruning processes that are shaped by both innate and postnatal factors. It was reported that human visual cortex thickens to a maximum during the first two years and gradually thins thereafter (Lyall et al., 2015), mirroring the trajectory of synapse in human striate cortex (Huttenlocher and Courten, 1987). The previous evidence that both congenitally blind and sighted subjects showed uptrend of CT in the early stage, leading to the conclusion that the visual experience might not be involved in this process (Bourgeois, 1996; Jiang et al., 2009). Our results in the human neonates suggested the postnatal experience could accelerate the thickening of neonatal visual cortex, even if it was not necessary.

Whether it is the visual experience out of various postnatal stimuli that modulated the CT of ventral visual cortex? Although we could not provide a direct evidence from the current study, there were indications that the CT in present study might reflect an experience-dependent synaptogenesis induced by early visual experience (Holtmaat and Svoboda, 2009). On the one hand, we found that across whole brain, CT increased most prominently in the V1, primary auditory area and central sulcus (Supplementary Fig 11), which directly receive visual, auditory, sensorimotor; while the increases of CM were mainly distributed in the frontal areas (Supplementary Fig 11). This whole brain pattern suggested that CT was shaped by postnatal environment universally and it is reasonable to expect that ventral visual cortex is closely related to visual experience. On the other hand, we found the hierarchically developmental pattern of cortical thickening in ventral cortex with the posterior area showing faster increase than anterior regions. This might reflect the influence of visual flow along the ventral cortex, e.g., the primary visual area in the posterior area processes all kinds of visual information while higher-level visual cortex responds less. Future study could combine the behavioral measurements to test the relation between this two factors or compare the sighted and congenitally blind infants to further illustrate such question.

T1w/T2w myelination measurement, which takes advantage of the covariation between cortical myelin content and T1w (positive covariation) as well as T2w (negative covariation) intensities, could be used as a surrogate maker of the cortical myelination (Glasser and van Essen, 2011; Soun et al., 2017). Cortical myelination is an important feature of neurodevelopment, and is related to various cognitive performance (Glasser et al., 2014). The increase of T1w/T2w intensity in visual cortex might reflect the augment of myelinated fibers in this area. Previous studies reported the coupling between cortical myelination and face processing, suggesting an experience-dependent development of visual cortical microstructure in later life (e.g. 5-12 years old; Nordt et al., 2021). We found that in the early stage, CM primarily correlated with the prenatal time (GA at birth) but not the extrinsic experience, suggesting that the early cortical myelination might be determined the endogenic growth. In addition, the correlation between CM and PMA was even stronger after we controlling the postnatal time, suggesting an inversed or insignificant effect of postnatal time on the speed of cortical myelination (Fig 2d, f). Such results might due to the change of nutrition environment, or the interaction between CT and CM, in which the rapidly thickening cortex diluted the increase of myelinated axons in specific cortex (Glasser and van Essen, 2011).

Using r-fMRI data, previous studies evaluated the functional organization of the visual system in neonatal macaques (Arcaro and Livingstone, 2017) and category-selective network in neonatal humans (6-57 days) (Kamps et al., 2020). Both two studies revealed the proto-organization prior to the emergence of functional domain such as face and place areas. Herein we validated those findings in a large sample with infants as early as one day of postnatal age, fully controlling the influence of postnatal visual experience on the results. Particularly, we found two distinct visual clusters along the medial-lateral axis, which captured the central-peripheral organization of ventral visual system (Grill-Spector and Weiner, 2014; Hasson et al., 2002; Levy et al., 2001; Wiesel, 1982). Apart from the border of two medial-lateral clusters, the lateral area become more face-selective while the medial area is specialized in processing scene and buildings later in life (Nordt et al., 2021), and thereby our observation suggests the innate scaffold is already established at birth for subsequent experience-dependent modification in ventral visual cortex (Arcaro et al., 2017; Arcaro and Livingstone, 2021, 2017). Furthermore, the connections among the ventral visual cortex have developed during this early stage. Specifically, the homotopic connections between bilateral V1 and between bilateral VOTC both increased with GA, indicating increased degree of functional distinction (Arcaro and Livingstone, 2017; Vincent et al., 2007); In contrast, the connection between V1 and VOTC decreased between 37 and 44 weeks of PMA, which might also reflect the development of functional differentiation between adjacent regions in the same hemisphere. More importantly, the connection between bilateral V1 were significantly modified by postnatal time, proving that the early postnatal experience not only influence the local structural features of cortical cortex but also the functional circuits. Similar to the finding in the structural development, the experience-dependent development of functional connectivity also followed a hierarchical manner, e.g., only the development of lower-level but not higher-level visual cortex showed postnatal dependence.

In brief, our results suggested that early cortical development is a mixed outcome of endogenous and experience-dependent development in both cortical structure and function, and experience could selectively modify the endogenous development of visual cortex during early infancy.

## Materials and Methods

### Participants

A total of 887 datasets (783 neonatal subjects) from developing human connectome project (dHCP: www.developingconnectome.org/) were collected. We excluded the datasets that 1) have high radiology score reviewed by specialist perinatal neuroradiologist (above 2 points), which indicated possibly clinical or analytical significance (e.g. punctate lesions or other focal white matter or cortical lesions but not consider to be of clinical significance; https://biomedia.github.io/dHCP-release-notes/structure.html) in the images (*n* = 218, including 2 datasets without scores); 2) were sedated during scan (*n* = 5); 3) were scanned early than 37 weeks (*n* = 91) of postmenstrual age (PMA); and 4) missing myelin maps or rest-state functional magnetic resonance imaging (r-fMRI) data (*n* = 152); 5) could not pass the quality control of dHCP preprocessing pipelines for the structural (*n* = 1) or r-fMRI data (*n* = 9). 6) failed the cortical registration pipeline (*n* = 4). Finally, 407 datasets (407 subjects) were selected into present study, in which 355 were term-born (GA: 39.93 ± 1.26 [37-42.29] weeks; PMA at scan: 41.14 ± 1.7 [37.43 – 44.71] weeks) and 52 were preterm-born (GA: 31.98 ± 3.35 [23.71-36.86] weeks; PMA at scan: 40.9 ± 1.91 [37 – 44.29] weeks). The dHCP study was approved by the UK Health Research Authority (14/LO/1169) and written consent was obtained from the parents or legal guardians of the study subjects.

### Data Acquisition

All neuroimaging data was acquired at the Evelina Newborn Imaging Centre, Evelina London Children’s Hospital using a 3-T Philips scanner with a newly designed neonatal imaging system including a customized 32-channel phased array head coil, an elaborated positioning device and a custom-made acoustic hood for infant (Hughes et al., 2017). Infants were imaged during natural sleep without sedation. Structural (T1 and T2 weighted image), rest-state functional (r-fMRI) and diffusion MRI (dMRI) were collected within a single scan session for each neonate over 63 minutes. T2-weighted (T2w) and inversion recovery T1-weighted (T1w) multi-slice fast spin-echo images were acquired in sagittal and axial stacks with in-plane resolution of 0.8 × 0.8 mm isotropic and slice thickness of 1.6 mm (0.8 mm overlap, except for sagittal T1w that used 0.74 mm). Other parameters were as follows: 1) T1w images were acquired with repetition time (TR) = 4795 ms, echo time (TE) = 8.7 ms, Inversion time (TI) = 1740 ms, SENSE factor of 2.27 (axial) and 2.66 (sagittal) and filed of view (FOV) = 145 × 122 × 100 mm; and 2) T2w images were acquired with TR = 12000 ms, TE = 156 ms, SENSE factor of 2.11 (axial) and 2.58 (sagittal) and FOV = 145 × 145 × 108 mm (the two acquired orthogonal stacks were integrated using a super-resolution method, Kuklisova-Murgasova et al., 2012). High temporal resolution r-fMRI optimized for neonates were collected echo-planar imaging using a multiband factor of 9 in 15.05 minutes: TR = 392 ms, TE = 38 ms, 2300 volumes, flip angle = 34° and spatial resolution = 2.15 mm isotropic (https://biomedia.github.io/dHCP-release-notes/acquire.html, Bozek et al., 2018; Fitzgibbon et al., 2020).

### Data Preprocessing

We collected the preprocessed anatomical and r-fMRI data from dHCP database (Makropoulos et al., 2018). For the anatomical data, briefly, preprocessing of dHCP pipeline included super-resolution reconstruction to obtain the 3D T1w/T2w volumes (Kuklisova-Murgasova et al., 2012), registration (from T1w to T2w), bias correction, brain extraction, segmentation (on T2w volume using DRAW-EM method, Makropoulos et al., 2014), surface extraction (Schuh et al., 2017), and surface registration (Robinson et al., 2018). The initial structural data from dHCP in present study included the individual brain surfaces, cortical thickness (CT, corrected version for uneven vertex sampling for gyri relative to sulci) and T1w/T2w myelination. Surfaces and cortical metrics of each neonate were nonlinearly aligned to the dHCP symmetric template (40 weeks of PMA; Bozek et al., 2018) using multimodal surface matching (MSM, Robinson et al., 2018). The transformation of 4 subjects failed so we exclude them from the following analysis.

The collected functional data from dHCP included the r-fMRI data in individual space and the motion parameters (Fitzgibbon et al., 2020). These images were further preprocessed with following steps using custom codes and DEPABI toolbox (Yan et al., 2016) in MATLAB (v2018a). 1) A conservative approach was adopted to address the severe head motion of infants. Similar to the previous study (Eyre et al., 2021), we selected a continuous subset (1600 volumes, around 70%) of the data with lowest head motion for each of the infants. Particularly, we first estimate the variation of head motion for each time point within a 50-volume window by calculating the standard deviation of the head motion within the time window, then we calculated the sum of the variations in all possible 1600 continuous volumes (e.g. 1-1600, 2-1601, …, 701-2300), and finally chose the subset with lowest sum of the head motion variation. 2) Registration of the selected r-fMRI subset to individual T2w space with FLIRT (Jenkinson et al., 2002). 3) Linear detrending. 4) Regression of nuisance covariates, which contained 24 head motion parameters, the signals of white matter, cerebrospinal fluid and global brain. These tissue masks were extracted from the T2w-based segmentation of the corresponding subject. 5) Temporal bandpass filtering with a pass band of 0.01 – 0.08 Hz. 6) The filtered r-fMRI volume was further projected into the individual cortical surface and then registered to the dHCP symmetric template using multimodal surface matching (MSM) method (Robinson et al., 2018). The resulting surface data was slightly smoothed with 2 mm full-width-half-maximum (FWHM) using workbench (Glasser et al., 2013).

### Anatomical ROIs

All ROIs were defined in adult space (Glasser et al., 2016), and registered into the dHCP template using MSM method with cortical sulcus as the features that drive the alignment (Robinson et al., 2018). The surface and sulcus data of adult atlas were the S1200 group-average data collected from the human connectome project (HCP; https://www.humanconnectome.org/). The congruent relationship of the parcellations between adult and infants was visually acceptable in large scales (Supplementary Fig 1-2) although the parcellation accuracy in refined areas could not be guaranteed.

The anatomical mask of ventral cortex was parcellated into 34 regions of interests (ROIs) per hemisphere in the HCP-MMP atlas (Glasser et al., 2016), which contain basic visual cortex (e.g. V1 and V2), higher-level visual cortex (e.g. ventral occipital-temporal cortex, VOTC) and anterior part of ventral temporal cortex (Supplementary Fig 1 and Supplementary Table 1). The VOTC includes 15 ROIs per hemisphere, including common category-selective regions such as parahippocampus, middle fusiform and lateral occipital cortex (Bi et al., 2016; Grill-Spector and Weiner, 2014). The primary visual area was defined as the V1 ROI in the HCP-MMP atlas (Supplementary Fig 1).

### Data Analysis

#### Experimental design

The purpose of this study is to describe the structural and functional development of ventral visual cortex in human newborns and evaluate the influence of postnatal experience in this process. To depict a comprehensive picture of the spatiotemporal development, for each structural or functional measurement, we first presented the general developmental trend of whole ventral cortex and also the spatial variations. Then we focused on the V1 and VOTC and characterized the detailed developmental pattern of these two areas. To evaluate the influence of postnatal experience on specific development, we first assessed Pearson correlation between individual measurements and postnatal time, and further used partial correlation to control the covariation between PMA and postnatal time in this dataset.

We further designed two sub-samples to validate the influence of postnatal effect in the development. The first sub-sample only includes infants who underwent the scans within 3 days after birth and thus limited postnatal experience, and the second sub-sample includes infants who had a limited PMA range from 40 to 42 weeks to estimate the effect of postnatal experience independent from PMA.

Lastly, we compared the term-born infants with preterm-born peers who had longer postnatal experience at equivalent PMA, which may help to explain the influence of early experience versus endogenous coding on cortical development. Therefore, for the structural and functional measurements which showed significant influence with postnatal time, we will further compare those measurements between term and preterm-born infants.

### Statistical analysis

For all of the correlation analysis, we removed the data beyond 3 SDs from the mean value across all infants. The correlation coefficients were Fisher-Z transformed before any statistical test, but the plot (e.g. box plot and scatter plot) used the original *r* value to give an intuitive picture. Extraction and calculation of cortical measurement (e.g. CT, CM and r-fMRI) were performed using gifti toolbox (https://www.nitrc.org/projects/gifti/) in MATLAB (v2018a) and commands in workbench (v1.5.0, https://github.com/Washington-University/workbench). Statistical analysis inlcuding Pearson correlation, partial correlation and *t* test were performed in the MATLAB. FDR correction (*q* < 0.05) were used for the multiple comparison correction if not specifically mentioned.

#### Structural data

##### Cortical thickness and myelination maps of ventral cortex between 38 to 44 weeks of PMA

We averaged cortical thickness and myelination map across neonates within each of the 9 PMA weeks (37 to 45 weeks). Because there were only 2 infants in the 37-week PMA group (37-37.5 weeks) and 1 infant in the 45-week PMA group (44.5 – 45.5), we did not present the averaged maps of those weeks.

##### Spatial differences between developmental trajectories

To quantify the spatial difference in the developmental change of these cortical measurements, we extracted the default coordinate system of the ventral cortex in the atlas, in which the y-axis was along the anterior-posterior direction and the x-axis was along the medial-lateral direction (Supplementary Fig 3a). For each axis, we divided the ventral cortex into 30 segments with equal length along the axis. For each segmentation, we averaged the correlation coefficients across all vertexes to obtain the spatial pattern of cortical development along the two axes (Supplementary Fig 3b-c).

##### Comparison of the developmental speed between different age groups in V1 and VOTC

We divided the subjects into two groups according to their age (PMA or postnatal age). The first group contain around 25% subjects with lowest age while the second group contain equal number of subjects with highest age. We used two-factors mixed ANOVA analysis with regions (V1 and VOTC) as the within-factor and neonatal age (high and low groups) as the between-factor to compare the age-dependency of cortical structural development (Fig 3d-e).

##### Degree of influence of prenatal time on cortical thickness and cortical myelination

We used two-factors mixed ANOVA analysis with cortical measurements as the within-factor (CT and CM) and neonatal prenatal time (subjects with highest and lowest 25% GA) as the between-factor (Supplementary Fig 3) to compare the influence of prenatal time on CT versus that on CM. We normalized the CT or CM to make them comparable, as

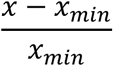

where *x* indicates the original CT or CM values, and *x*_*min*_ indicates the minimum value in the group.

#### R-fMRI data

##### Temporal correlation analysis

Pearson correlation were carried out on the averaged time series (across all vertex in the 34 ROIs) within and across left and right ventral cortex. The arealization for each of the 34 area was calculated using Pearson correlation between the contralateral (homotopic) ROIs in two hemispheres, and the *p* values were corrected with Bonferroni method (corrected *p* < 0.01). To compare the correlation between homotopic ROIs and non-homotopic ROIs in the ventral cortex, we applied two-sample *t* test between 34 homotopic connections and the rest of the non-homotopic connections (e.g. V1-V2, V2-V3 and so on).

##### Multi-dimensional scaling

Similar to the method used in the previous studies (Arcaro and Livingstone, 2017), for each hemisphere, we obtained the ipsilateral correlation matrix between the 34 patches in ventral cortex. Then the correlation matrix was transformed into a distance matrix using 1 – *r* for each entry of the matrix. Non-classical multi-dimensional scaling (MDS) was carried on the distance matrix with Kruskal’s normalized criterion in MATLAB. The 34 ROIs were plotted in a two-dimension coordinate according to the first two principal values in the MDS analysis.

##### Clustering analysis

For the correlation matrix between the 34 Rois within ipsilateral ventral cortex, a threshold of *r* = 0.15 was used to remove weak and negative connections and regional clusters in the ventral cortical network were obtained using a spectral algorithm (Newman, 2006) in the Brain Connectivity Toolbox (BCT; Rubinov and Sporns, 2010). Different threshold (e.g. 0.1 and 0.2) were also used to validate the results (Supplementary Fig 6).

##### Developmental changes of ventral network

Two typical network measurements including global efficiency and mean cluster coefficient (calculated in BCT) were used to quantify the integration and segregation of the ventral network. Pearson correlation between two measurements and PMA (or postnatal time) was used to describe the development of the ventral network in human newborns.

##### Developmental change of the connections in V1 and VOTC

Four ROIs including V1 and VOTC in two hemispheres were included in this analysis. Pearson correlation was conducted between the averaged time series in each 4 ROIs, resulting in two contralateral connections between V1 and VOTC, and two ipsilateral connections between V1 and VOTC in each hemisphere that were then averaged for each subject. We calculated the Pearson correlation coefficient between these connections and PMA (or postnatal time) to measure the developmental changes.

#### Comparison between term-born and preterm infants

Due to the unbalanced sample size between preterm-born (*n* = 52) and term-born infants (*n* = 355), we selected a sub-group in the term-born infants with equal sample size and similar PMA (40.92 ± 1.87 weeks) to the preterm-born neonates (40.90 ± 1.91 weeks; *t* = 0.06, *p* > 0.95). Particularly, for each infant in the preterm group, we chose a non-repetitive subject in the term-born whose PMA was closest to the preterm-born infant. Structural and functional measurements were compared between two groups using two sample *t* test.

## Data availability

The data used in this work are available online on the dHCP page (http://www.developingconnectome.org/data-release/third-data-release/).

## Code availability

Custom-written MATLAB codes used for data preprocessing and analysis are available upon reasonable request.

## Acknowledgements

This work supported by the Ministry of Science and Technology of the People’s Republic of China (2018YFE0114600), National Natural Science Foundation of China (61801424, 81971606, 82122032), and Science and Technology Department of Zhejiang Province (202006140). Data were provided by the developing Human Connectome Project, KCL-Imperial-Oxford Consortium funded by the European Research Council under the European Union Seventh Framework Programme (FP/2007-2013) / ERC Grant Agreement no. 319456. We are grateful to the families who generously supported this trial.

## Author contributions

M.L., X.D. and D.W. conceived this work. M.L. performed all the data analysis, with contributions from T.L., X.X., Q.W., Z.Z., X.D., Y.Z. and D.W. The manuscript was written by M.L. and D.W. and all the authors edited it.

## Competing interests

The authors declare no competing interests.

## Notes

### Competing Interest Statement

The authors have declared no competing interest.

