## Supplemental figures and table for "Development of visual cortex in human neonates are selectively modified by postnatal experience"

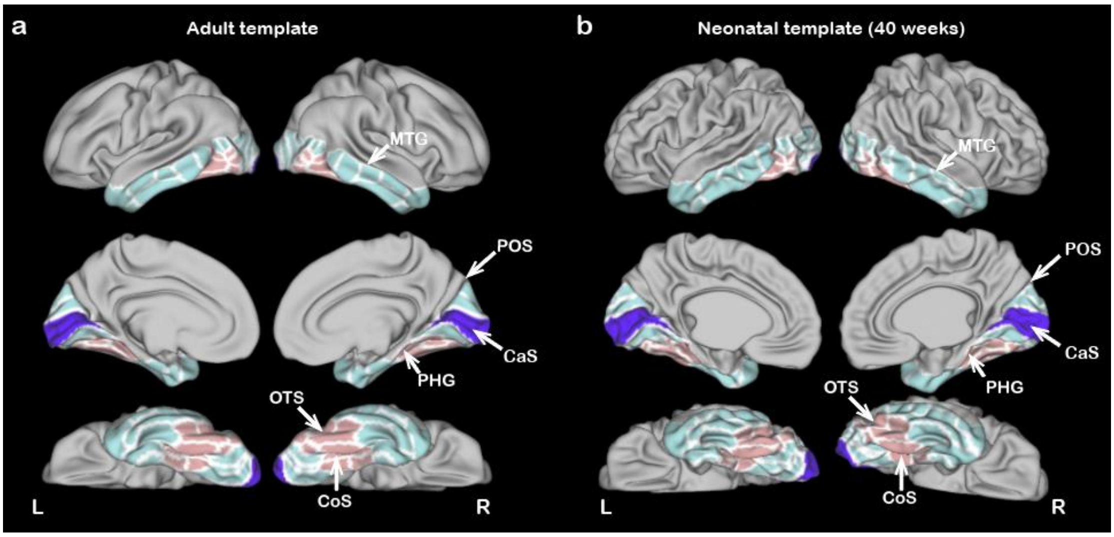


**Supplementary Fig 1.** The mask of ventral cortex was parcellated into 34 ROIs in HCP-MMP atlas (a), which was projected into neonatal template at 40 weeks of PMA (b). White arrows indicated the homologous landmarks in the two atlases: 1) The parietooccipital sulcus (POS) was the posterior-upper boundary of the mask; 2) the middle temporal gyrus (MTG) and parahippocampal gyrus (PHG) served the lateral and medial boundary of the mask; 3) the calcarine sulcus (CaS) was occupied by the primary visual cortex (V1, purple regions); and 4) the occipitotemporal sulcus (OTS) and the collateral sulcus (CoS) were included in the higher-level visual cortex (pink regions).

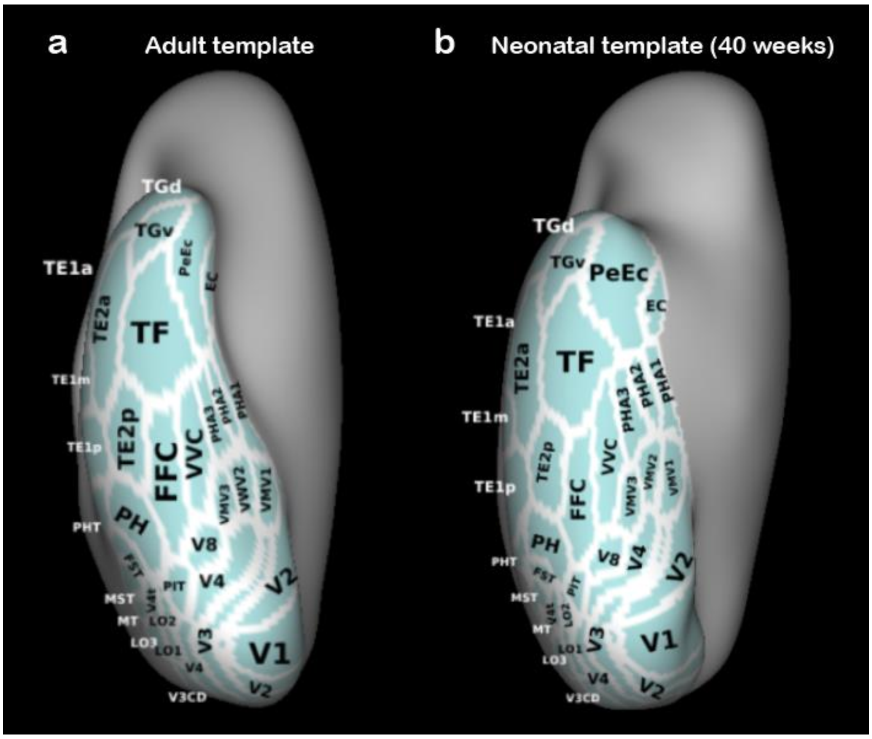


**Supplementary Fig 2.** The detailed labels of each ROI in the ventral view of adult (a) and neonatal (b) very-inflated atlas in the right hemisphere. The area descriptions are provided in the Supplementary Table 1.

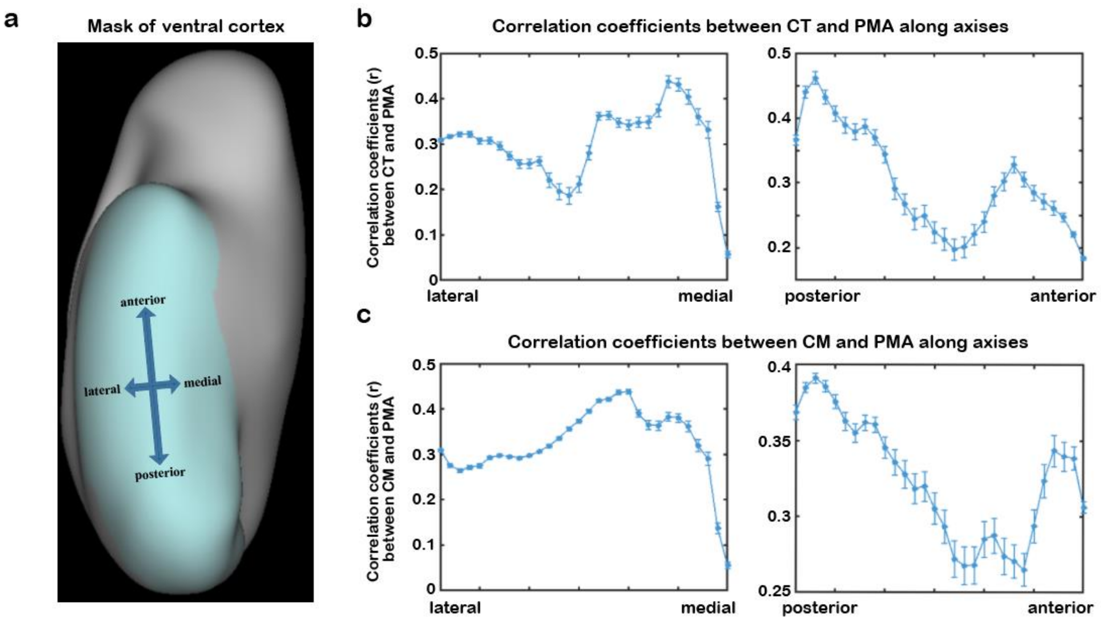


**Supplementary Fig 3.** To quantify the spatial variation of the development of (b) CT and (c) CM from 37 to 44 weeks of PMA (Fig 2a), we divide the ventral cortex into 30 segments with equal length along anterior-posterior or medial-lateral axes (a). The correlation coefficients between PMA and CT/CM were averaged across all vertexes in each segments and plotted along the two axes. Error bar indicates the standard error within the segment.

Note: PMA = postmenstrual age; CT = cortical thickness; CM = cortical myelination.

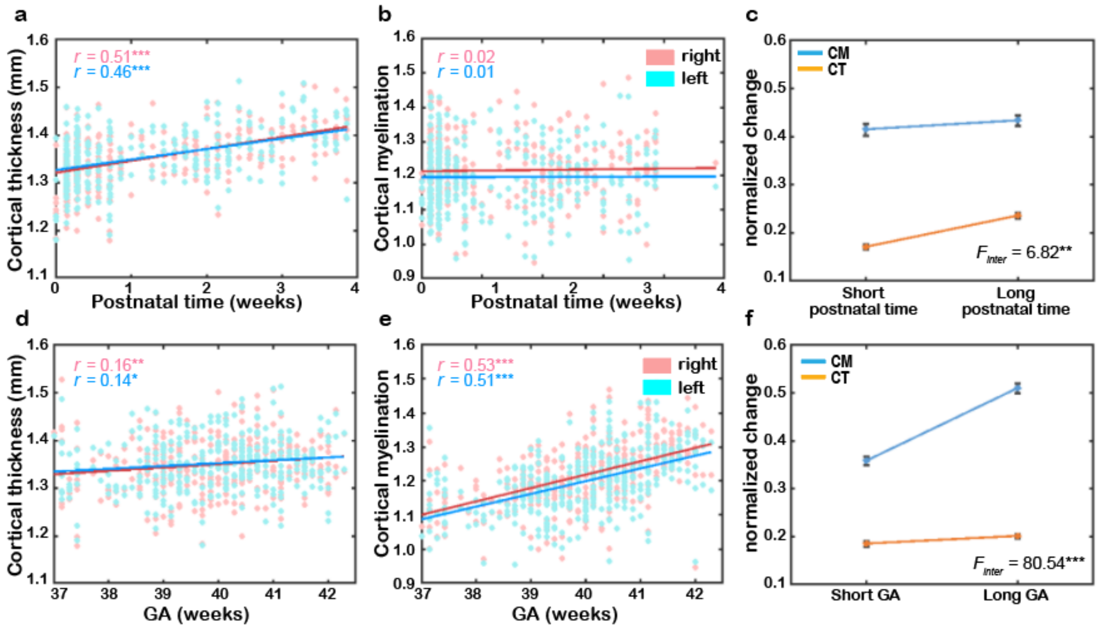


**Supplementary Fig 4.** The correlation between postnatal time or prenatal time (GA at birth) and CT or CM (a,b,d,e), indicating that CT was heavily dependent on the postnatal time but the CM was dependent on the prenatal time. The interaction between the normalized change of CT (orange) and CM (blue) in the short and long postnatal (c) (or prenatal; f) time groups further supported that the CT and CM were differently influenced by postnatal and prenatal time.

Note: GA = gestational age; CT = cortical thickness; CM = cortical myelination; $F_{\text{inter}}$ = the $F$ value of the interaction between the time (short and long postnatal or prenatal group) and the normalized changes of structural measurements (CM vs. CT) using two factor mixed ANOVA analysis. * $p < 0.05$, ** $p < 0.01$, *** $p < 0.001$.

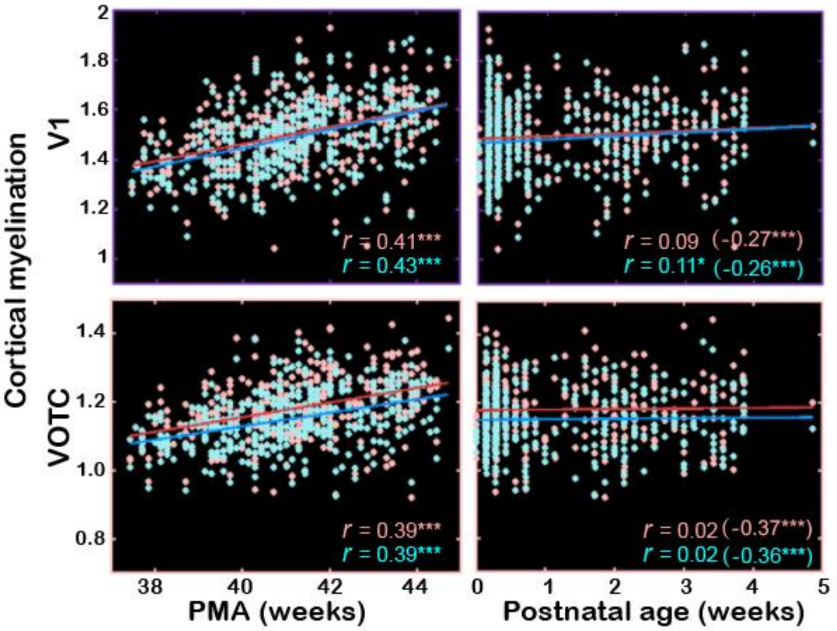


**Supplementary Fig 5.** Correlation between cortical myelination and PMA or postnatal age in V1 and VOTC. The values in the bracket indicate the partial correlation coefficients between CM and postnatal time when controlling for PMA.

Note: PMA = postmenstrual age; CM = cortical myelination; V1 = primary visual cortex; VOTC = ventral occipital temporal cortex.

* $p < 0.05$; ** $p < 0.01$ and *** $p < 0.001$.

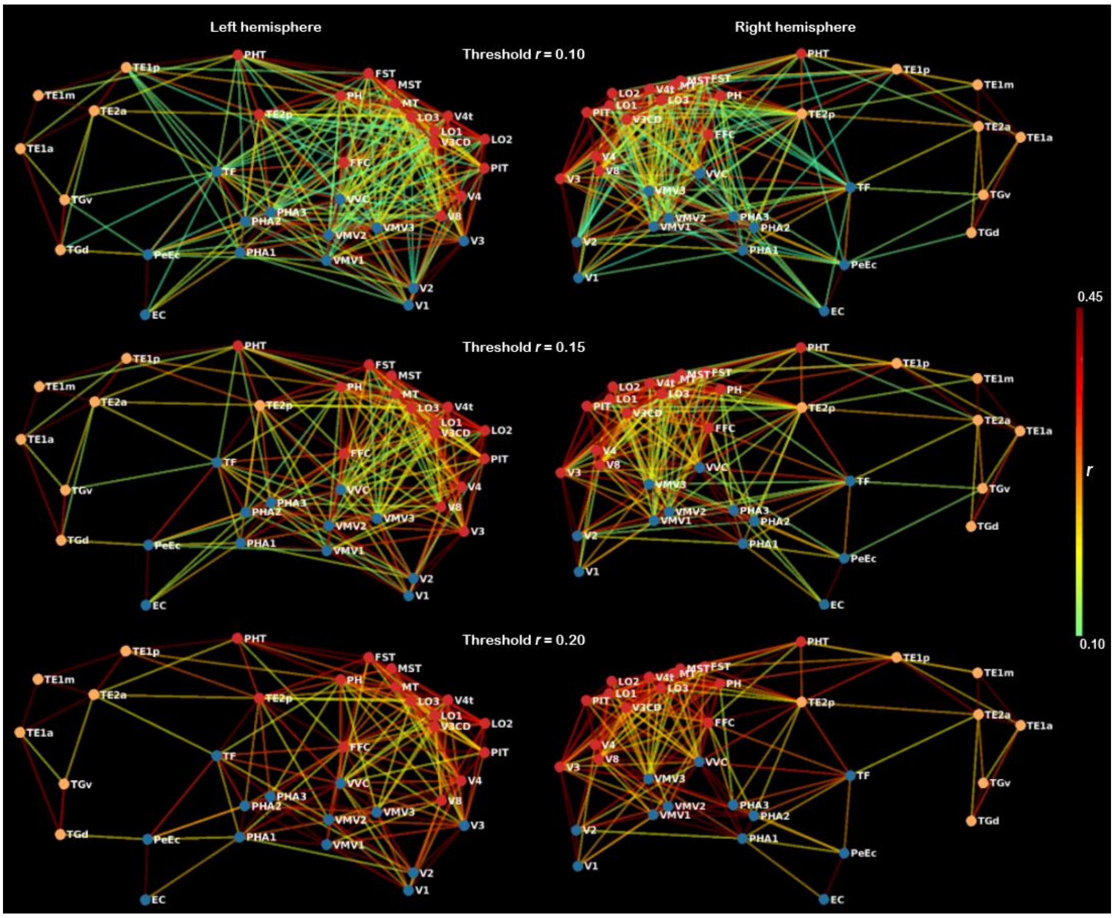


**Supplementary Fig 6.** Multidimensional scaling and community groups based on pairwise ipsilateral connections between 34 ROIs in the left or right ventral cortex. The results were similar between left and right cortex, and similar in the networks using different thresholds of correlation coefficient *r*. The color of the nodes indicates the cluster identity of each node in the community structure analysis and color of the lines connecting the nodes indicates the functional correlation *r* between two nodes within the same hemisphere.

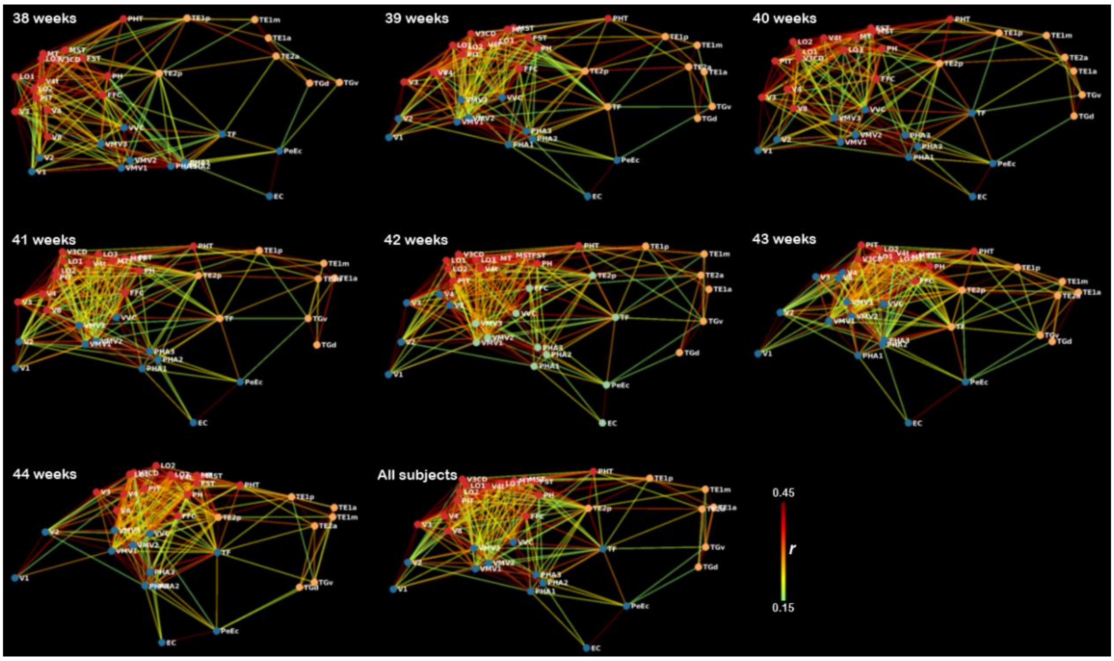


**Supplementary Fig 7.** Functional networks of right ventral cortex for each PMA week from 38-44 weeks, based on multidimensional scaling. We found similar three-cluster network structure in the infants at different PMA (except for 42 weeks), and subtle fluctuation was observed in the boundary between different clusters at different PMA.

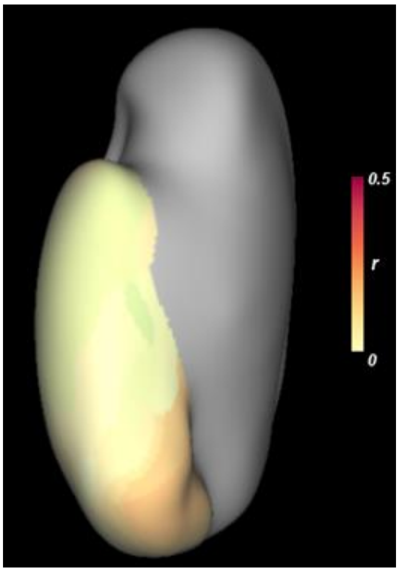


**Supplementary Fig 8.** The partial correlation between homotopic connections and postnatal age controlling for PMA of infants.

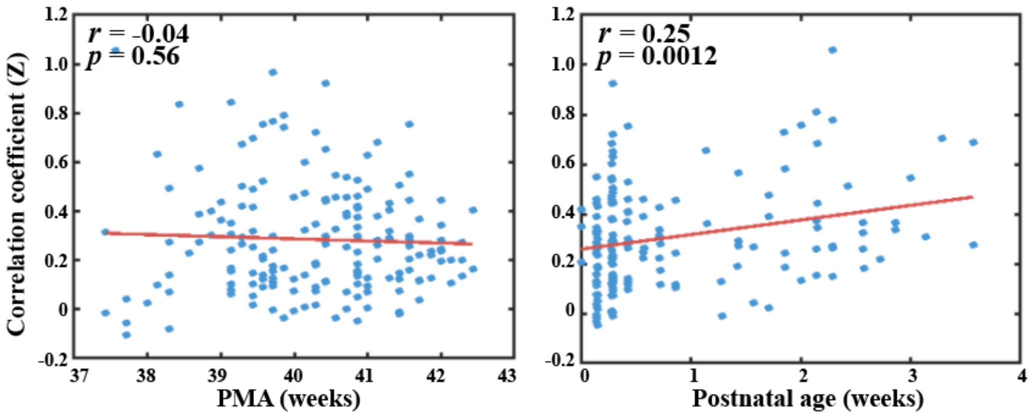


**Supplementary Fig 9.** The correlations between bilateral V1 connection and PMA (left) or postnatal age (right) in two sub-samples, indicating that the postnatal experience could increase the functional connection between bilateral V1, and such effect was probably independent of the PMA. The y-axis indicate the Fisher-Z transformed *r* value.

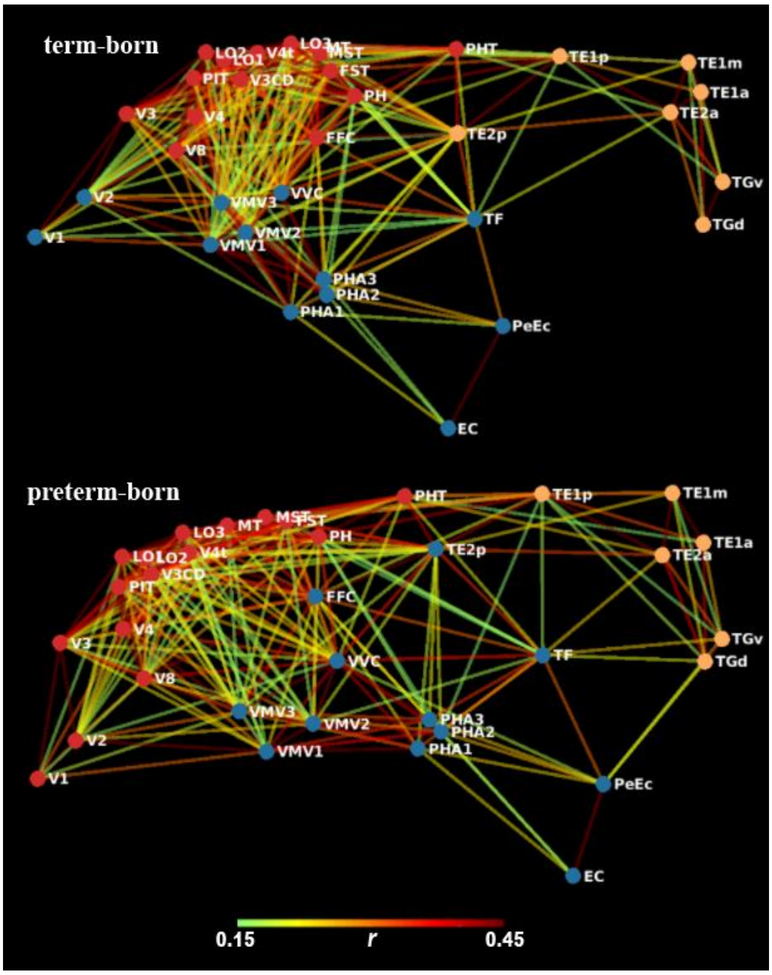


**Supplementary Fig 10.** Functional networks of right ventral cortex in term-born and preterm-born infants, based on multidimensional analysis.

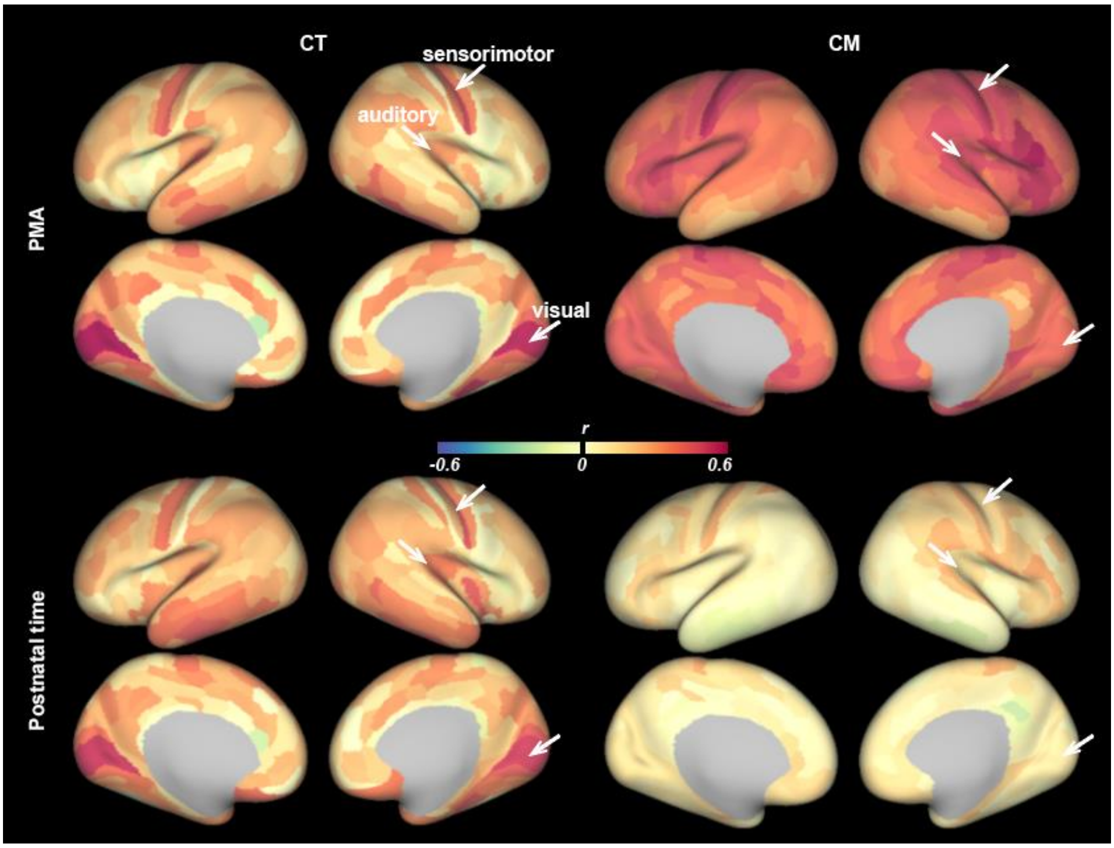


**Supplementary Fig 11.** The correlation coefficient maps between PMA or postnatal age and the structural measurements (CT or CM) across the whole brain. The primary sensory areas such as V1 (visual), primary auditory area (auditory) and central sulcus (sensorimotor) demonstrated the most prominent increase of CT with respect to both PMA and postnatal time, suggesting the universal effect of postnatal sensory experience on the development of CT.

Note: PMA = postmenstrual age; CT = cortical thickness; CM = cortical myelination.

**Supplementary Table 1.** The labels of 34 ROIs in the ventral cortex and 15 of the 34 ROIs (bolded) are in the ventral occipital temporal cortex.

| **Number** | **Area Name** | **Area Description** |
|---|---|---|
| 1 | V1 | Primary Visual Cortex |
| 2 | V2 | Second Visual Area |
| 3 | V3 | Third Visual Area |
| 4 | V4 | Fourth Visual Area |
| **5** | **V8** | **Eighth Visual Area** |
| 6 | V3CD | Area V3CD |
| 7 | LO3 | Area Lateral Occipital 3 |
| 8 | LO1 | AreaLateral Occipital 1 |
| 9 | MT | Middle Temporal Area |
| 10 | MST | Medial Superior Temporal Area |
| **11** | **V4t** | **Area V4t** |
| **12** | **LO2** | **Area Lateral Occipital 2** |
| **13** | **FST** | **Area FST** |
| **14** | **PIT** | **Posterior InferoTemporal complex** |
| **15** | **PH** | **Area PH** |
| **16** | **TE2p** | **Area TE2 posterior** |
| **17** | **FFC** | **Fusiform Face Complex** |
| **18** | **VVC** | **Ventral Visual Complex** |
| **19** | **VMV3** | **VentroMedial Visual Area3** |
| **20** | **VMV2** | **VentroMedial Visual Area 2** |
| **21** | **VMV1** | **VentroMedial Visual Area 1** |
| **22** | **PHA3** | **ParaHippocampal Area 3** |
| **23** | **PHA2** | **ParaHippocampal Area 2** |
| **24** | **PHA1** | **ParaHippocampal Area 1** |
| 25 | PHT | Area PHT |
| 26 | TE1p | Area TE1 posterior |
| 27 | TE1m | Area TE1 middle |
| 28 | TE1a | Area TE1 anterior |
| 29 | TGd | Area TG dorsal |
| 30 | TE2a | Area TE2 anterior |
| 31 | TF | Area TF |
| 32 | EC | Entorhinal Cortex |
| 33 | PeEc | Perirhinal Ectorhinal Cortex |
| 34 | TGv | Area TG Ventral |

Note: the area names and area descriptions were identical to the original literature.
